# Versatile Encapsulation and Synthesis of Potent Therapeutic Liposomes by Thermal Equilibration

**DOI:** 10.1101/2021.10.22.465473

**Authors:** Steven A. Roberts, Chaebin Lee, Shrishti Singh, Nitin Agrawal

**Author notes:** Corresponding Author, 111 Michigan Ave NW, Washington, DC 20010, USA. Equal Contribution (Chaebin Lee, Shrishti Singh).

## Abstract

The wide-scale use of liposomal delivery systems is hampered by difficulties in obtaining potent liposomal suspensions. Passive and active loading strategies have been proposed to formulate drug encapsulated liposomes, but are limited by low efficiencies (passive) or high drug specificities (active). Here, we present an efficient and universal loading strategy for synthesizing therapeutic liposomes. Integrating a thermal equilibration technique with our unique liposome synthesis approach, co-loaded targeting liposomes can be engineered in an efficient and scalable manner with potencies 200-fold higher than typical passive encapsulation techniques. We demonstrate this capability through simultaneous co-loading of hydrophilic and hydrophobic small molecules and through targeted delivery of liposomal Doxorubicin to a metastatic breast cancer cell line MDA-MB-231. Molecular dynamic simulations are used to explain interactions between Doxorubicin and liposome membrane during thermal equilibration. By addressing the existing challenges, we have developed an unparalleled approach that will facilitate the formulation of novel theranostic and pharmaceutical strategies.

## Introduction

Decades of research have resulted in a variety of liposomal formulations for use in both therapeutic^1–3^ and theranostic^4–6^ applications. Advances in synthesis procedure and membrane functionalization have led to targeting and stealth capabilities, which can increase bioavailability at disease sites several fold with minimal side effects, compared to systemically administered drugs^7–8^. These improvements have in turn led to the FDA approval and commercial use of several liposomal drugs, including Doxil^®^ for cancer^9^, Abelcet^®^ for fungal infections^10^, and DepoDur^®^ for pain management^11^, and more recent Moderna/Pfizer vaccines for Covid-19^12^. While substantial progress has been made in the development of novel liposomal formulations, considerably fewer strategies to increase the potency and concentration of therapeutics within the vehicles have been developed.

Sequestration of therapeutic compounds has traditionally been accomplished by either passive or active encapsulation approaches. The passive approach can sequester molecules irrespective of their polarity^13^, where lipophilic molecules reside within the lipid bilayer membrane and hydrophilic molecules are contained within the core. During vesiculation, liposomes form within the media containing drug, encapsulating the drug in the process. This approach is typically done with dilute solutions, and therefore the encapsulation efficiencies are relatively low ^14^. This leaves much of the drug to be lost during secondary purification steps. Furthermore, organic solvents (e.g. chloroform and ethanol) are typically used, that can inactivate certain molecules^15^.

Active loading techniques have been developed that drastically improve encapsulation efficiencies and minimize the amount of non-encapsulated drug. The sequestration is driven by transmembrane pH or ion gradients that cause the influx of hydrophilic molecules to the core of liposomes^16–18^. This approach has been successfully utilized to sequester weakly basic drugs with encapsulation efficiencies of nearly 100%. Once inside the liposome, drug molecules precipitate out of the solution, increasing their retention time. While this technique overcomes many of the barriers to passive encapsulation, it is limited to the molecules that can pass through the membrane. Furthermore, decorating the membrane with targeting moieties may require a second step to remove any unbound targeting molecules that may result in the loss of encapsulated drugs.

We have previously demonstrated a simple approach for producing purified drug loaded liposomes^19^. Using this method, we rapidly synthesize concentrated populations of small unilamellar liposomes (SULs) with a very narrow polydispersity index (typically near 0.13). The approach relies on passive encapsulation to sequester small molecules, and as such, the encapsulation efficiency is relatively low. Therefore, a technique that can enhance the encapsulation efficiency utilizing small amount of compounds is desired. Here, we demonstrate a new approach for creating drug-loaded targeting liposomes using thermal equilibration, where drug molecules are sequestered at high concentrations via passive diffusion at the membrane transition temperature (T_m_). We demonstrate the universal and translational appeal of this approach by simultaneously co-loading both hydrophilic and lipophilic small molecules. We further demonstrate the therapeutic potential of this technique by encapsulating Doxorubicin (DXR), without utilizing ion gradients typically required for active methods. Bioactivity of these liposomes was observed by actively targeting cancer cells *in vitro*. Finally, using molecular dynamic simulations, we explain the thermodynamics that underlie the encapsulation and model the effects of temperature on structural properties of the liposome membrane and DXR diffusion. The thermal equilibration approach presented here is a powerful universal strategy for rapidly encapsulating small molecule drugs. It offers significantly greater encapsulation capabilities than conventional passive encapsulation techniques and represents a feasible solution to producing therapeutic grade liposomes in a scalable manner.

## Experimental

### Materials

Cholesterol, 1,2-Distearoyl-sn-glycero-3-phosphocholine (DSPC), 5,5′-Dithiobis(2-nitrobenzoic acid) (Ellan’s reagent), L-Cysteine, and all fluorescent dyes were purchased from Sigma Aldrich (St. Louis, MO). 1,2-distearoyl-sn-glycero-3-phosphoethanolamine-N-[maleimide(polyethylene glycol)-2000] (ammonium salt) (DSPE-PEG-Mal) was purchased from Avanti Polar lipids. Isopropyl Alcohol (IPA, 99% Pure), Ethanol (EtOH, 99% pure), Dimethyl Sulfoxide (DMSO), desalting columns, dye removal columns, and Trypsin-EDTA were purchased from Fisher Scientific (Hampton, NH). Glass syringes were purchased from Hamilton (Reno, NV) and luer lock dispensing needles were purchased from Jensen Global (Santa Barbara, CA). Amicon Ultra 100 kDa centrifugal filter units were purchased from EMD-Millipore (Billerica, MA). A NanoJet syringe pump from Chemyx Inc. (Stafford, TX) was used for all experiments. Dulbecco’s Modified Eagle Medium (DMEM), Fetal Bovine Serum (FBS), and Penicillin-Streptomycin antibiotic were purchased from VWR (Radnor, PA). Doxorubicin (DXR) and WST-1 proliferation assay were purchased from Cayman Chemical (Ann Arbor, Michigan). MDA-MB-231 metastatic breast cancer cell line was acquired from ATCC (Manassas, VA).

### Synthesis of Small Unilamellar Liposomes

Liposomes were synthesized as previously described^19^. Briefly, DSPC and Cholesterol were dissolved in isopropyl alcohol at a constant 2:1 molar ratio to a final concentration of 10 mM and 5 mM, respectively. The alcoholic solution was injected using a syringe pump at 100 μL · min^-1^ into a glass vial containing distilled water preheated at 55°C to obtain a homogeneous population of liposomes. Typical synthesis volumes would be a total of 10 mL (1 mL alcoholic lipid suspension to 9 mL warm distilled water). The aqueous solution was constantly stirred at 600 rpm during infusion, and stirring was continued for 3 minutes following infusion. After the liposomal suspension was cooled down to room temperature with continuous stirring, liposomes were concentrated 20 fold to a volume of 500 μL and a final lipid concentration of 20 mM using filter centrifugation (100 kDa, 6000 x g) at 4°C for 30 minutes. The particles were stored at 4°C until use.

### Antibody Coupling

Antibody coupling was accomplished through the addition of functionalized PEGylated lipids (DSPE-PEG-Mal). DSPE-PEG-Mal was dissolved to a concentration of 25mg/mL in DMSO and added to the liposomal suspension containing small molecules (e.g. DXR, Nile Red, or Fluorescein) to a final molar ratio of 1% (DSPE/DSPC). The suspension was incubated for one hour at 55°C to increase fluidity of the membrane. Simultaneously, antibodies targeting CD44 were thiolated by incubation with Traut’s reagent at a ratio of 3:1 (protein: Traut) for 1 hour. Unreacted Traut’s reagent was removed by passing antibody suspensions through a desalting column and the resulting thiolated antibody was added to the liposomal suspension at a molar ratio of 1:1 (DSPE:Protein). The coupling reaction occurred overnight at 4°C. Finally, non-reacted maleimide groups were neutralized by a 15 minute incubation with β-mercaptoethanol at a molar concentration of 3:1 (β-mercaptoethanol:DSPE). The resulting suspension was then filtered as previously described to remove all non-encapsulated and unreacted molecules.

### Quantification of PEG-maleimide and Coupling Efficiency on the Liposomes

The amount of DSPE-PEG-maleimide on liposomes was evaluated following Ellman’s test^20^. Briefly, 120 μL of the functionalized liposomes were mixed with 30 μL of L-Cysteine (0.36 mM) using a thermomixer at 700 rpm for 2 hours at room temperature. The particles were removed by filter centrifugation for 30 minutes at 6000 x g. In a transparent 96-well plate, 100 μL of the supernatant and 50 μL of 5,5′-Dithiobis(2-nitrobenzoic acid) (0.144 mM) were added and mixed for 30 minutes at room temperature. The absorbance at 409 nm was measured using a microplate reader, and the unreacted L-Cysteine was determined using a standard curve. The amount of conjugated DSPE-PEG-maleimide was calculated by subtracting unreacted L-Cysteine from the total amount of L-Cysteine added to the functionalized liposomes, and the DSPE-PEG-MAL to DSPC ratio was evaluated.

### Thermal Equilibration for Small Molecule Encapsulation

In this work, a hydrophilic (Fluorescein sodium salt) and two hydrophobic compounds (Nile red and Doxorubicin) were selected for encapsulation using thermal equilibration. Fluorescein sodium salt was prepared in water at a concentration of 1.5 M while Nile Red and Doxorubicin HCl were prepared in DMSO at concentrations of 30 mM and 172 mM, respectively. The concentrations of small molecule are detailed in the results section. The small molecule solution was added to liposome suspension and incubated at desired temperatures and times. After liposomes were cooled to 4°C, non-encapsulated DXR was removed from suspension using a Zeba Spin desalting column with a 7 kDa cutoff, while excess fluorescent dyes were removed using dye removal columns. For targeted liposomes, DSPE-PEG-Mal was dissolved to a concentration of 25 mg/mL in DMSO and added to a final mol ratio of 1% (DSPE/DSPC).

### Analysis of Small Molecule Encapsulation and Release

Encapsulated compound concentrations were analyzed using a Thermo Scientific Varioskan Flash multimode reader. DXR concentration was quantified using an excitation of 525 nm and an emission of 580 nm. Liposomes were solubilized in 0.1% Triton x-100 before measuring the fluorescence. The fluorescence intensity was compared to the standard curve which was prepared by measuring the fluorescence of DXR in 0.1% Triton x-100. To quantify the concentration of co-loaded samples, after dissolving liposomes in 0.1% Triton x-100, the fluorescence was measured at an excitation of 488 nm and emission of 512 nm in water for fluorescein. The samples were then desiccated for several hours under vacuum and redissolved in IPA. The fluorescence of Nile Red was measured using an excitation of 550 nm and an emission of 625 nm. The fluorescence intensities of fluorescein and Nile Red from the sample were compared to standard curves prepared with either triton x-100 in water (fluorescein) or IPA (Nile Red) to evaluate encapsulated compound concentrations.

DXR release was investigated by diluting samples 20-fold in PBS with 10% FBS and incubating for up to 48 hours at 37°C. At indicated time points, liposome suspension was purified by multiple filtrations to remove released DXR. First, the liposome suspension was filtered through a 7 kDa desalting column to remove free doxorubicin followed by another filtration using a 100 kDa Amicon ultra centrifugal filter unit to remove any serum bound DXR. Liposomes were dissolved in 0.1% Triton x-100, and the fluorescence of each sample was measured using an excitation of 525 nm and an emission of 580 nm. The remaining DXR in liposomes was calculated by comparing the fluorescence intensities to a standard curve prepared with DXR in 0.1% triton x-100.

### Stability of Liposomes in Serum

The effect of serum on liposome stability was tested by monitoring the particle size change in either H2O, PBS, or PBS with 10% FBS for up to 24 hours at 37°C. Briefly, 25 μL of liposome solution was dispersed into either H2O, PBS, or PBS with 10% FBS and mixed by thermomixer at 37°C. Samples were collected at different time points (0, 1, 4, 16, and 24 hours), and 20 μL of each sample was introduced into 500 μL of fresh H2O for size measurement using DLS.

### Quantification of Phospholipid Concentration

Phospholipid concentration was empirically determined via Stewart Assay^21^. Briefly, liposomal suspensions were dehydrated at 70°C in polypropylene microcentrifuge tubes. The lipids were then dissolved in 500uL chloroform, and an equal volume of Stewart’s reagent (100 mM FeCl_3_ and 400 mM NH_4_SCN in DI water) was added and vortexed for 20 seconds. The tubes were centrifuged for 10 minutes at 1000 x g to create a phase separation. The chloroform layer was then analyzed in a quartz cuvette for its absorbance at 472 nm and compared to a standard curve to determine the phospholipid concentration. The standard curve was made using a serial dilution of lipids in cholesterol at a 2:1 ratio.

### Cell Culture and Therapeutic Delivery

MDA-MB-231 cells were grown in DMEM containing 10% FBS and 1%Penicillin-Streptomycin solution. The cells were cultured at 70% confluence and plated in a 96 well plate at a density of 5000 cells/well. The cells were incubated undisturbed for two days prior to any experimentation. Liposomal suspensions were UV sterilized prior to introduction to cell cultures. On the second day, the liposomal carriers containing DXR were introduced to the cultures at the IC_50_ of DXR (0.01 μM^22^). Proliferating cells were quantified and normalized by the control using WST-1 proliferation assay following the manufactural protocol.

### Membrane Simulations

Simulations are performed on a section of the lipid membrane consisting of DSPC and cholesterol held in a 2:1 molar ratio. The membrane section was constructed using Charmm-27 GUI Membrane Builder^23^, and designed to consist of 256 total lipid molecules and 128 cholesterol molecules. The membrane was hydrated using 30 water molecules per lipid. Nanoscale Molecular Dynamics (NAMD) was used to thermally equilibrate the bilayer system for 1 ns at designated temperatures. Production runs were also conducted at designated temperatures with a time step of 1 femtosecond and data collection every 20 femtoseconds. Every simulation was run for at least 100 ns, yielding at least 5000 data points per temperature. Trajectories of the simulations were visualized using Visual Molecular Dynamics (VMD). Further information on the simulation setup is available in the supplementary information.

### Statistical Data Analysis

Statistical analysis was conducted using GraphPad Prism 5 (La Jolla, CA). For each condition, at least three samples were independently prepared, and each sample was analyzed three times. Statistical analysis was performed using either a student’s unpaired t-test or a One-Way Analysis of Variance (ANOVA) with a Tukey post-test. Data was deemed statistically significant if p values were less than 0.05. Graphs show the mean and the standard error of the mean (SEM) of sample groups.

## Results

### Synthesis Approach and Maximum Equilibration Calculation

Typical liposome synthesis techniques^15,24,25^ require extensive secondary processing (e.g. sonication^26^, extrusion^27^ freeze thawing^28^) to generate monodisperse SULs. An alternative method that has been utilized to generate SUL’s without extensive post processing is through the injection method (**Figure 1a**). The injection method is advantageous in its simplicity but typically yields low encapsulation efficiencies^19^. To address this, we have developed a novel approach to efficiently and rapidly sequester small molecules.

**Figure 1:**
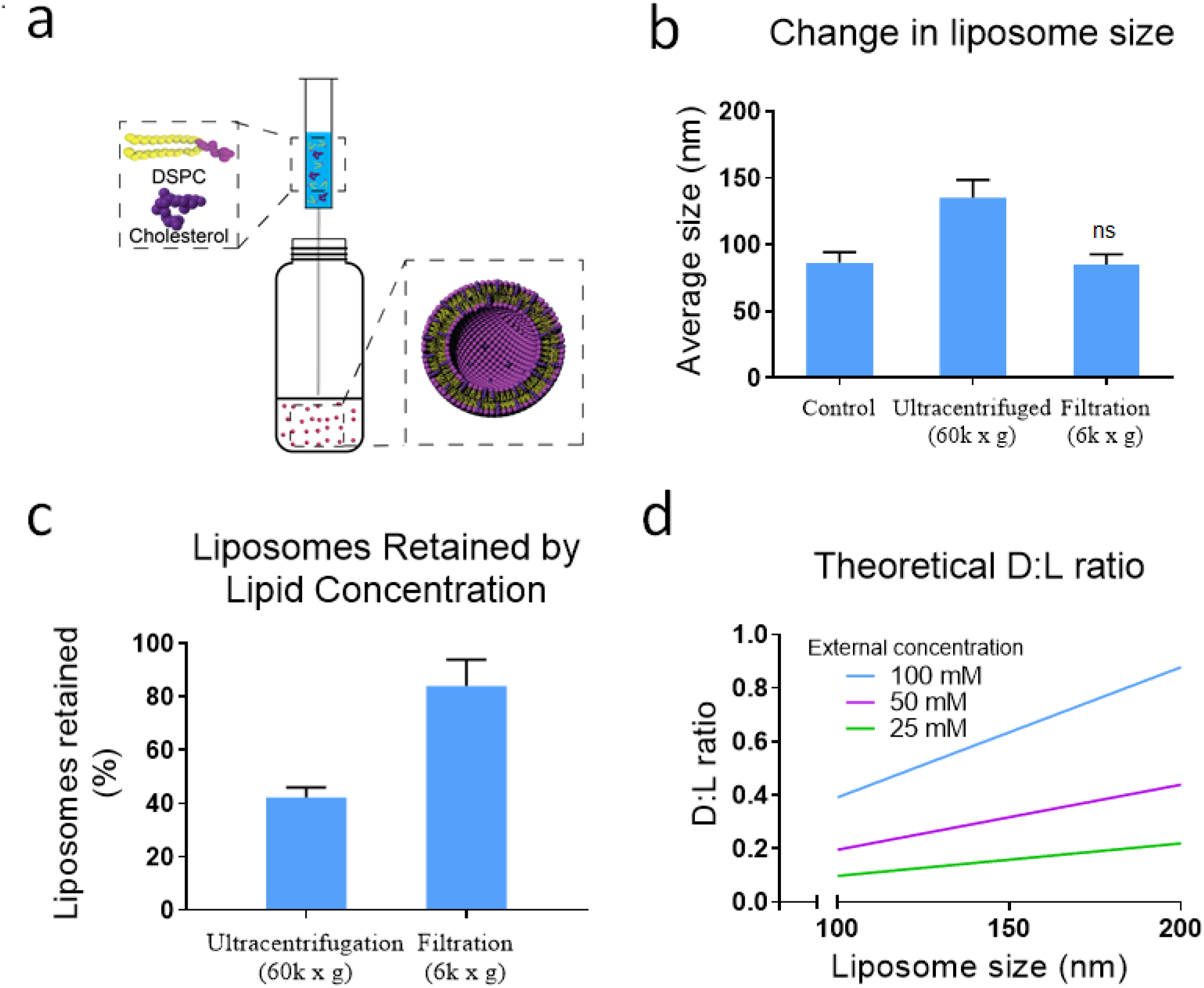
Synthesis of potent targeted liposomes. (a) Liposomes are synthesized in a rapid and scalable manner using the injection method. Membrane components are dissolved in isopropyl alcohol and injected into DI water at 55°C. The phase change causes the arrangement of liposome components to form small unilamellar liposomes. The solution is concentrated 20x using filter centrifugation. (b and c) The filtration process retains the population of liposomes efficiently, as seen in the size distribution and lipid concentration. The liposome population increases in diameter from 86.6 nm to 135.4 nm when ultracentrifuged as a result of small liposomes remaining in the supernatant. The change in diameter when filtered was not statistically significant, suggesting the relative population was retained. Analysis of lipid concentration showed nearly 60% of the liposomes were lost following ultracentrifugation, whereas filtration retained 84% of the total liposomes in solution. (d) The theoretical D:L ratio of thermal equilibration for hydrophilic molecules is dependent on liposome diameter and initial drug concentration, where increasing liposome diameter and external concentration results in an increase in liposome potency.

#### Liposome Purification and Concentration

Small unilamellar liposomes are difficult to concentrate without losing part of the population in the process due to their low densities^29,30^. Therefore, we proposed to use filter centrifugation instead of ultracentrifugation. To compare the efficiency of ultracentrifugation and filter centrifugation, raw liposome without loading any compound were tested. Empty liposomes were homogeneous with a polydispersity of less than 0.2. By filter centrifuging for thirty minutes at 6000 x g, we effectively concentrated the liposomes 20-fold while retaining the characteristic size and lipid concentration **(Figure 1b and 1c)**. The mean population diameter of liposomes increased from 86.6 nm (± 4.5 nm) following synthesis to 135.3 nm (± 7.7 nm) following ultracentrifugation at 60000 x g for two hours. Furthermore, by analyzing the lipid concentration, we found that only 42% (± 2.1%) of the lipids were retained, suggesting that ultracentrifugation can only pellet larger liposomes within the population. As further proof of this, we analyzed supernatant of the ultracentrifuged sample using DLS and routinely found liposomes with diameters of 80 nm (data not shown). Notably, there was a non-significant change in liposome diameter following filter centrifugation (85.3 ± 4.3 nm), and 79.2% (±1.3%) of the lipids were retained. While increased centrifugation speeds (>60,000 x g) and longer centrifugation durations using a specialized ultracentrifugation system may improve liposome retention, the proposed filter centrifugation approach is significantly more efficient and accessible.

#### Measurement and Quantification of Drug Encapsulation

We evaluated the maximum equilibration concentration (C_max_), where the number of drug molecules within the liposomes (C_E_) in solution will be proportional to the number of drug molecules in the surrounding media (C_T_). The constant of proportionality (α) here is the ratio of the respective volumes, where V_Excluded_ is the cumulative core volume of all liposomes and V_Total_ is the total volume of the solution.

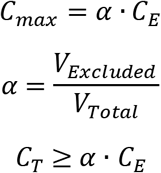

V_Excluded_ can be estimated by determining the inner volume of one liposome and extrapolating for the entire liposome population within the sample. Furthermore, if the drug is lipophilic, V_Excluded_ can be substituted for the volume of the lipid membrane. Using our maximum equilibration model, we can predict the D:L ratio as a linear function of concentration and liposome size (**Figure 1d**). This function is linear due to the constant lipid molarity, meaning as the liposome size grows, it utilizes lipids that could otherwise be used to form additional liposomes. Additionally, if the density of the lipid membrane does not change as liposome size changes, the maximum equilibration of lipophilic drugs is solely dependent on C_T_. Therefore, when equilibrating liposomes with a narrow size distribution, we can use C_max_ to hypothesize the liposome potency based on extra-liposomal drug concentration (dependent on drug solubility) as well as the number of liposomes present. By concentrating the samples using filter centrifugation, we can increase the C_max_ and therefore the potency of liposomes.

#### Stability of Liposomes during Thermal Equilibration

Several factors can affect the diffusion of small molecules across the membrane, including temperature of the system and solvents used. Alcohols such as isopropanol and ethanol offer convenient methods to increase the solubility of many drugs, however, these solvents can also deactivate or precipitate some drugs. DMSO is an alternative that is widely used to enhance the solubility of nonpolar chemicals for drug delivery. To investigate the effect of organic solvent on liposomal stability particularly at temperatures near T_m_, we individually incubated aqueous solutions of isopropanol, ethanol, and DMSO with concentrated liposomal slurries at 55°C and analyzed their size change. We found that 10% DMSO had the least effect on liposome size (**Figure 2a**), with an average shift from 104 nm (± 0.35 nm) to 107 nm (± 0.55 nm). Membrane components immediately precipitated out of the solution when introduced to alcohol concentrations of greater than 40%. Samples that include 10% DMSO were stable for at least 24 hours at 55°C (**Figure 2b**). In all cases, we found that the stability of liposomes diminishes rapidly when exposed to temperatures above T_m_.

**Figure 2:**
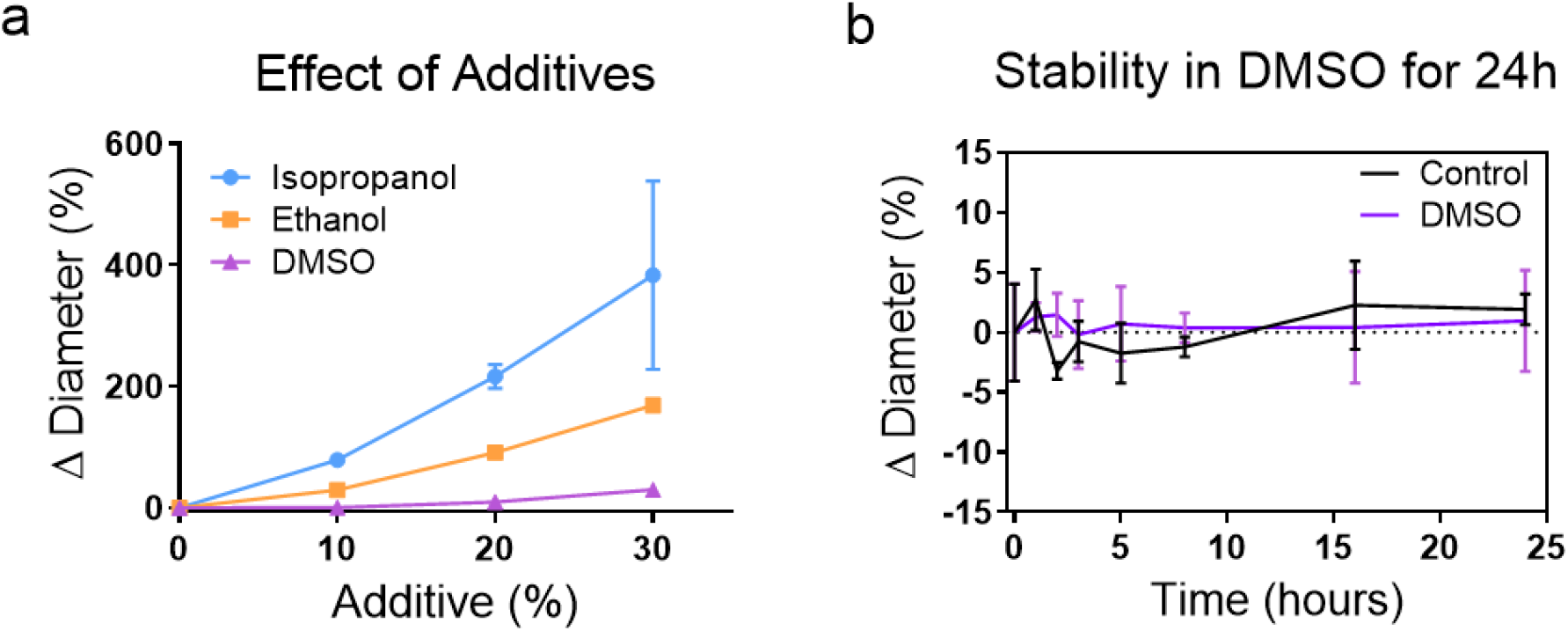
Analyzing the effect of temperature and solvents on liposome stability. (a) Liposomes incubated at 55°C for one hour with 10% DMSO did not show a statistically significant change in mean diameter (p=0.9565). However higher levels of DMSO and all tested concentrations of IPA and EtOH drastically affected the population. (b) Liposomes incubated at 55°C with and without 10% DMSO were stable for up to 24 hours and showed no significant change in liposome diameter.

### Encapsulation and Delivery of Chemotherapeutics from Equilibrated Liposomes

#### Equilibrating Liposomes with Doxorubicin

After characterizing the stability of liposomes during thermal equilibration, we evaluated the liposomal sequestration of therapeutic compounds. Thermal equilibration (**Figure 3a**) relies on the incubation of concentrated liposomal solutions, where the ratio of drug:lipid (D:L) can be optimized to enhance the potency of the vehicle without using large amounts of therapeutic compounds. Liposomes were incubated with DXR being 10% of the final concentration (D:L ratios of liposomal suspension were at 1.72:1). To validate our hypothesis of temperature affecting thermal equilibration, liposomal slurries were incubated with DXR at varying temperatures ranging from 4 to 55°C (**Figure 3b**). We also compared this approach with another commonly used approach of passive encapsulation^14^. When held at identical initial D:L levels, equilibration at T_m_ produced nearly 200 times higher D:L ratios compared to passive encapsulation (0.000239 and 0.041, respectively). While this is lower than current active forms of doxorubicin sequestration, which can achieve D:L of nearly 0.3, it is also less prohibitive and can be applicable to a larger library of molecules. Interestingly, we found that a significant amount of DXR had associated with liposomes equilibrated below T_m_ (D:L ratio of 0.011), which we believe may be due to advantageous binding. We observed a clear exponential trend with time during the initial one-hour incubation (**Figure 3c**). We found no difference in liposome diameter following thermal equilibration with DXR solubilized in DMSO (**Supplementary Figure 1**).

**Figure 3:**
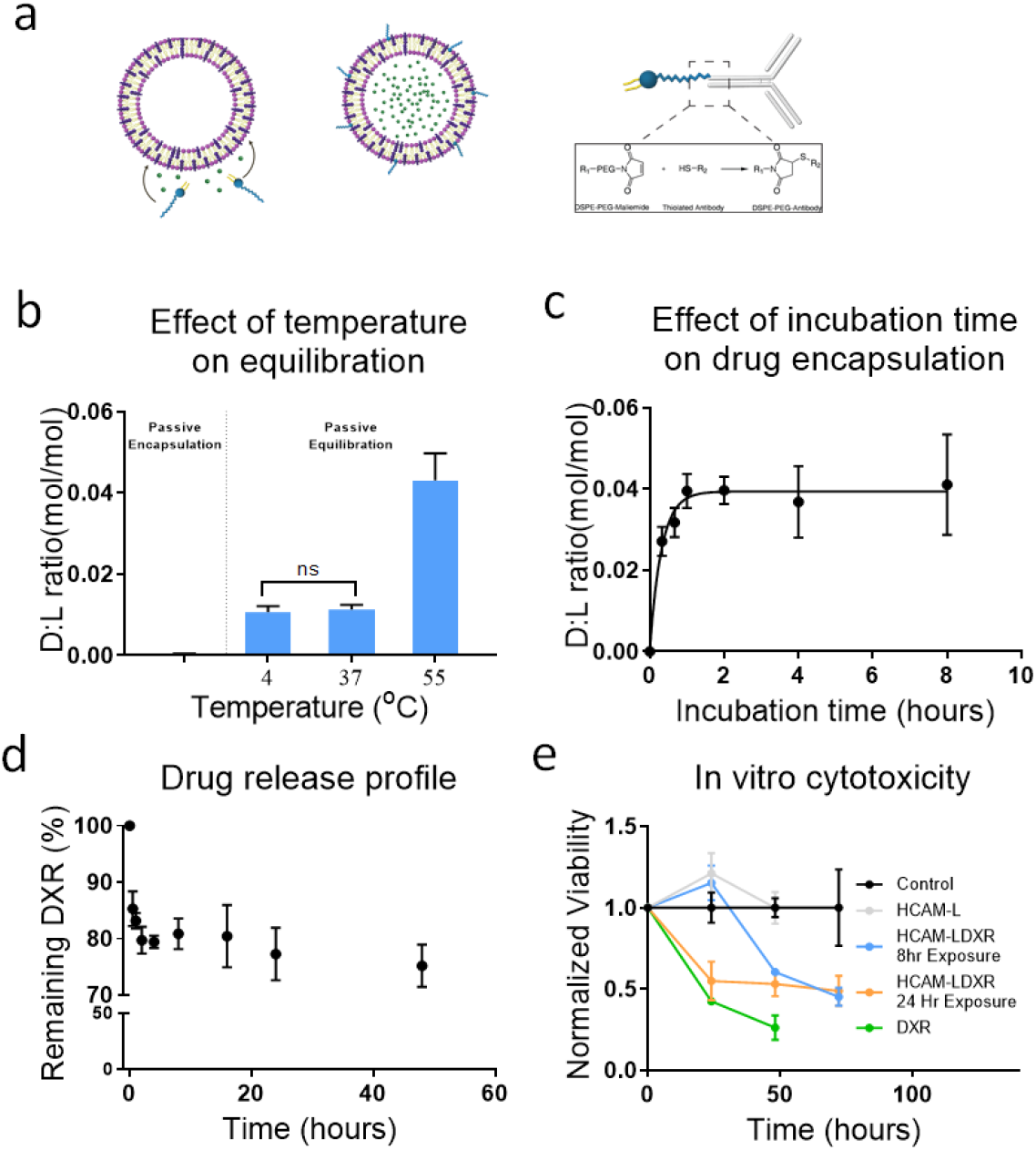
Thermal equilibration (a) Liposomes are equilibrated by incubating concentrated slurries with small molecules. Simultaneously, the liposomes can be incubated with DSPE-PEG-MAL linked to an antibody. This pegylated form of the potent liposome can be coupled to antibodies for efficient drug delivery. (b) The effect of temperature on equilibration was investigated by incubating DXR and liposomes at 4, 37, and 55°C for one hour. These results were compared to liposomal DXR created via passive encapsulation. E_T_ increases D:L nearly 200 folds compared to passive encapsulation, and even at low temperatures DXR is associated with the liposomes. (c) To optimize the encapsulation approach, liposomal samples were equilibrated for varying durations (up to 8 hours) with DXR. At indicated time intervals, liposomes were purified and the amount of DXR entrapped was measured through spectrometry. The liposomes showed a time dependent increase in DXR encapsulation up to 1 hour, however beyond 1 hour there was very little change. (d) Evaluations of the release of DXR were conducted by incubating E_T_ liposomal DXR at 37°C in PBS with 10% FBS. The results demonstrate that there is a controlled escape of DXR from liposomes, with 75% (±37%) of the DXR remaining in the liposomes after 48 hours. (e) During equilibration, DSPE-PEG-MAL can be integrated within the membrane, which when incubated with an antibody (HCAM), generates potent targeted liposomes. These targeted liposomes were cytotoxic when introduced to cancer cells *in vitro*, with a 52% decrease in cancer cell viability following 3 days of incubation. All measurements on the encapsulated DXR concentrations were determined using spectrometry and comparing absorbance values to a standard curve of known concentrations. Data was taken on at least 3 independently produced samples (n=3) and measurements on each sample were made in triplicate.

A hallmark of drug delivery systems such as liposomes is their stability and slow release of therapeutics under physiological conditions. This is especially important for the delivery of chemotherapeutic compounds such as DXR which can have severe toxic side effects^31^. Furthermore, extravasation into thick tumors may require 24-48 hours^32–34^, and therefore formulations that release compounds too quickly are not capable of reaching the required therapeutic index upon reaching the target site. Furthermore, interaction with serum is known to induce liposome degradation. We found that the liposomes remain stable in water and PBS without change in size at least for 24 hours at 37°C, whereas an increase in size of approximately 10% from 105.9 ± 10.2 nm to 117.8 ± 15.3 nm was observed when incubated with 10% FBS. The stability of thermally equilibrated liposomes under 10% FBS condition was confirmed by slow release of DXR with 75% of the encapsulated compound being retained after 48 hours (**Figure 3d, Supplementary Figure 2**).

#### Delivery and Cytotoxicity of Doxorubicin Liposomes

A distinguishing characteristic of drug delivery systems is the capability to provide therapeutic benefits through cellular uptake. To demonstrate this, we created drug loaded liposomes using the thermal equilibration technique. Liposomes were incubated with DXR (17.2mM) and DSPE-PEG-MAL (1 mol%) for 1 hour^35,36^ at 55°C followed by purification using spin columns. However, it is likely that part of DSPE-PEG-MAL will remain free, forming micelles that would be difficult to filter and subsequently interfere with antibody coupling. Therefore, retention of DSPE-PEG-MAL was calculated by determining the ratio of DSPE-PEG-MAL to DSPC lipid by Ellman’s test (**Supplementary Figure 3**). The DSPE-PEG-MAL to DSPC ratio of 0.009 ± 0.001 was observed, as compared to the starting ratio of 0.01 based on 1 mol% concentration suggesting that >90% of DSPE-PEG-MAL was retained within the solution following the insertion protocol. The subsequent CD44 conjugation and cancer cell targeting with fluorescein loaded liposomes displayed a strong signal of the liposomal uptake further validating that neither the DSPE-PEG-MAL molecules nor the CD44 antibodies were lost as micelles or during filtration (**Figure 5c**).

CD44 is highly expressed on MDA-MB-231 cells^37^ and was thus used to target DSPE-PEG-MAL incorporated liposomes after functionalization with the thiolated anti-CD44 to the maleimide group. Targeting liposomal DXR (LDXR) was incubated with MDA-MB-231 (at IC_50_) for 8 or 24 hours prior to being washed away with PBS. To ensure DMSO did not play a role in cell death, vehicles were generated by incubating pre-formed liposomes with DMSO in the absence of DXR. These liposomes were also functionalized with anti-CD44 IgG (HCAM-L). Free DXR was used as a positive control. Cell viability was analyzed over 72 hours using WST-1 as an indicator of proliferation and normalized to the media only control (**Figure 3e**). Cells demonstrated a time dependent decline in proliferation, as expected, when incubated with any form of DXR. Both 8 and 24 hours LDXR exposures eventually ended with nearly equivalent (p=0.173 at 48 hours and p=0.603 at 72 hours, n>3) viabilities.

### Atomic Level Simulation of Membrane Dynamics

Based on the predicted maximum equilibration ratios, liposomes with an average size of 100 nm, should have D:L ratios of 0.4 (**Figure 1d**). The prediction assumes that the inner core of the liposome contains the same concentration of compound as the external environment at equilibrium. Since the encapsulation curtails at 1 hour (**Figure 3c**), the liposomes are assumed near equilibrated by then, with D:L ratios at 68% of the maximum equilibrium concentration after which it plateaued. We were interested in what might be causing the difference between predicted and observed D:L ratios, and what forces cause the association of DXR with the liposome membrane at temperatures below T_m_. We hypothesized that the observed association between DXR and liposomes at temperatures <T_m_ may be due to the interactions between the phosphate group in DSPC and NH_2_ group of DXR. Using molecular dynamic simulations, we investigated the molecular interactions at the atomic level.

Since the passage of DXR is largely mediated by diffusion, movement through the membrane is dictated by both the long range Van Der Waals (VdW) energy and short range electrostatic forces existing between DXR and lipid molecules^38^ (**Figure 4a and 4b**, respectively). We explored the interactions, the NH_2_ group of DXR has with both the head and tail of the lipid. The analysis of VdW indicated that at 37°C, the DXR molecule primarily interacts with the lipid head (average energy −43.8 kcal/mol), while interactions between DXR and the tails are negligible (average energy −1.9 kcal/mol). Similarly, the electrostatic interactions at the head (**Figure 4b**) are highly attractive promoting the observed adsorption at temperatures lower than T_m_ (**Figure 3b**). Conversely, at T_m_ (55°C) when the lipid layer is more fluidic, the DXR first contacts the head (as indicated by the low VdW), but also moves closer to the tails (as evident by the decreasing VdW). At and above the T_m_, as the DXR moves into the membrane, and thus further away from the charged phosphate heads, the attraction decreases (KdW becomes more positive), before returning to a more negative steady state. Interactions between the leaflet and the DXR molecule are illustrated in **Figure 4c**. Below T_m_, DXR remains within the Van der Waals radii, interacting with the phosphate heads. However, as the membrane acquires a more liquid-state, the lipophilic region of DXR migrates into the tails. Most of the inter-atomic interactions occur within 5-8 Å of each other (**Figure 4d**), suggesting weak transient interactions, with the majority occurring between DXR and the lipid head.

**Figure 4:**
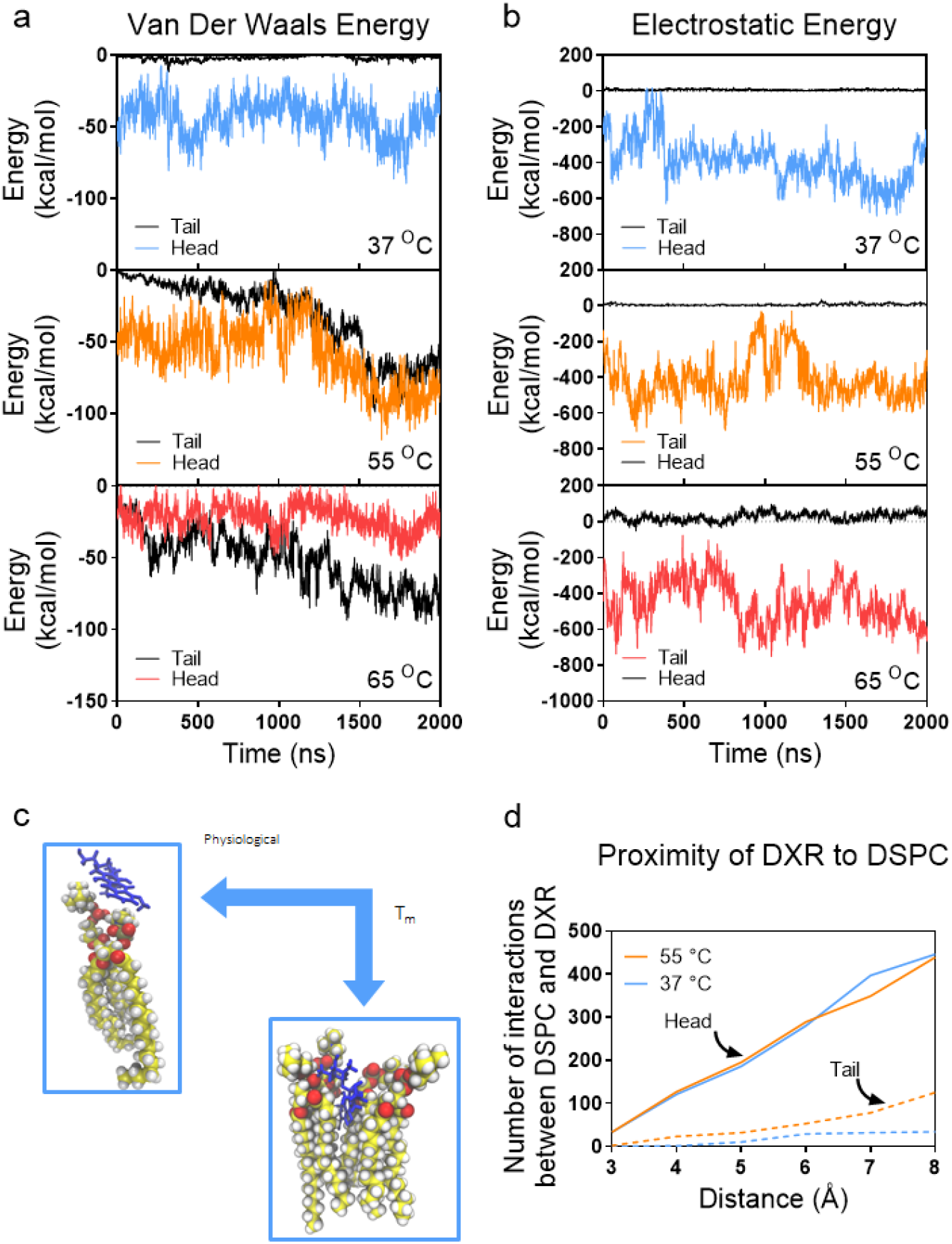
Atomic level simulations. Analysis of (a) long-range VdW and (b) short range electrostatic forces. At all temperatures, the DXR molecules are attracted to the negatively charged lipid heads, however only as the membrane acquires a more liquid-state (≥T_m_) does the DXR shows any attraction or interaction to the lipid tails. (c) The simulations depict an adsorption of DXR to the membrane below T_m_, however at and above T_m_ DXR infiltrates into the membrane. (d) Most DXR atoms are located within 5-8 Å of DSPC, insinuating a more additive and transient non-covalent bond.

While temperature clearly plays a role in thermal equilibration of liposomes with their exterior, experimentally we observed an almost immediate deleterious response in liposome stability at temperatures greater than T_m_. To elucidate this at the atomic level, we explored the effects of temperature on other factors that can influence DXR diffusion and membrane stabilization (**Supplementary Figure 4a**). In addition to the electrostatic and VdW interactions, the DXR molecule can stabilize via the additive effect of hydrogen bonding (**Supplementary Figure 4b**). While the number of bonds initially starts equally, irrespective of temperature, there is a significant decrease as the temperature increases and as the DXR enters the membrane. Conversely, at lower temperatures, the hydrogen bonding steadily increases, thereby stabilizing the interactions. Interestingly, while there is a significant decrease in the average number of hydrogen bonds when increasing the temperature from 37°C to 55°C (12.75 vs 6.75, respectively) there is very little difference between the number of hydrogen bonds at 55°C and 65°C (averages 6.75 and 4.5, respectively). This decrease in hydrogen bonding at temperatures ≥T_m_ likely enhances the DXR’s ability to migrate into the tail section, however, the levels are re-established as the DXR progresses through the membrane, suggesting a rotation of the DXR molecule so that the NH_2_ group points towards the polar heads. These results are most evident in the RMSD of the DXR, where there is an increase in the average DXR movement of over 50% at T_m_ compared to 37°C, but only an 8% increase between 55°C and 65°C (**Supplementary Figure 4c**). Therefore, while there is a large difference in equilibration between 37°C and 55°C, the change is not proportional above T_m_.

The membrane fluidity that permits DXR infiltration, as well as the loss of stability that correlates with the observed liposome disintegration at higher temperatures, is seen most apparently in the conformational energy of the tails (**Supplementary Figure 4d**). While higher conformational energy (and thus membrane fluidity) will enhance DXR infiltration into the membrane^39^, our observations noted that liposomes rapidly destabilize at temperatures > T_m_. Over the 18°C difference between gel phase (37°C) and the T_m_ (55°C), the conformational energy is increased 2.51%. Whereas, a smaller 10°C change in the temperature from 55°C to 65°C netted a similar 2.52% increase in conformational energy. Taken together with our experimental results, we believe that this non-proportional increase in conformational energy results in the observed decrease in liposome stability. While liposomes can mitigate the initial increase in conformational energy, for at least our observation window (24 hours) (**Figure 2c and 3b**), they are unable to remain stable with the additional increase in energy. Therefore, while increasing the temperatures beyond T_m_ may increase DXR movement (**Figure 4c**), it also greatly decreases the membrane stability and thus does not offer a feasible approach to increase equilibration.

### The Universal Nature of the Thermal Equilibration Approach

Many popular encapsulation approaches are drug specific and are thus not versatile. However, we foresee using passive equilibration as a universal encapsulation technique that could be applicable to a variety of molecules. To demonstrate this, we chose two small molecules that vastly vary in their polarity, Fluorescein, and Nile Red. These molecules are similar in size and have compatible excitation and emission spectra, which allows us to view the co-loading capabilities of our approach (**Figure 5a**). Fluorescein and Nile Red (solubilized in DMSO) were introduced to liposomal suspensions at concentrations of 750 mM and 20 mM, respectively. Samples showed D:L levels of 0.03 (±0.013) and 0.003 (±0.000099), respectively (**Figure 5b**).

**Figure 5:**
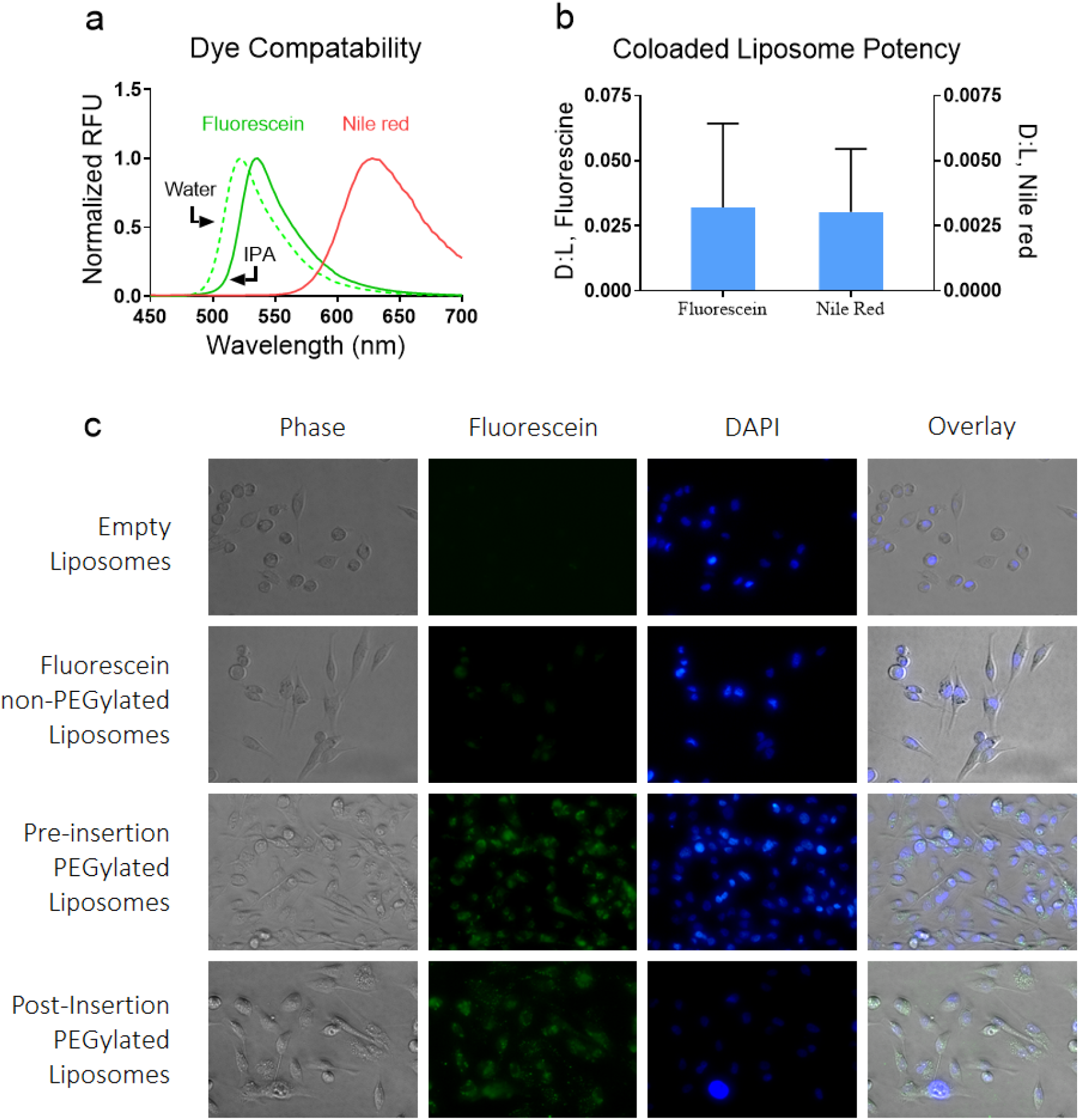
The use of thermal equilibration as a universal encapsulation method. Liposomes are uniquely capable of sequestering both lipophilic and hydrophilic molecules. (a) Using the equilibration approach, liposomes were loaded with the hydrophilic and lipophilic dyes Fluorescein and Nile Red, respectively. These dyes differ greatly in fluorescence emission and can be easily distinguished. (b) The liposomes were then co-loaded by incubating with a mix of fluorescein and Nile Red. Analysis of liposome emission spectra reveal two peaks indicative of both dyes being sequestered. (c) Liposomes loaded with fluorescein (green) were targeted to MDA-MB-231 breast cancer cells in vitro using PEG-DSPE-Mal covalently bonded to CD44 as a targeting molecule. The PEGylated lipid was introduced either during vesiculation or during thermal equilibration.

Furthermore, we demonstrate efficient targeting of liposomes to cancer cells and *in vitro* cellular uptake using the proposed technique (**Figure 5c**). Here, PEGylated liposomes were created through “Pre-insertion” (DSPE-PEG-Mal was added into the IPA solution pre-synthesis), or “Post-insertion” (DSPE-PEG-Mal was integrated in following the synthesis via thermal equilibration along with fluorescein). Both approaches yielded functionalized targeting liposomes, as demonstrated by the presence of fluorescein when examining cells exposed to PEGylated liposomes, and comparing their fluorescence to cells introduced to non-PEGylated liposomes.

## Discussion

We show here a universal approach to sequester small molecules within liposomes for drug delivery, irrespective of the drug polarity. We hypothesized that by incubating liposomes with small volumes of high concentration drug solution, the internal environment of liposomes would eventually equilibrate with the external environment, with localization largely depending on the partitioning coefficient of the compound (**Figure 3a**). We further hypothesized that this effect could be enhanced by increasing the fluidity of the membrane (via thermal incubation), which would also allow the introduction of targeting moieties within the membrane.

A lipid vesicle’s T_m_ is a physical characteristic that dictates the order of the lipid membrane at varying temperatures, and it is a crucial variable to consider in engineering liposomal delivery systems, where synthetic lipids have a higher T_m_ than natural lipids^40^. DSPC is a synthetic, saturated, 18 carbon lipid with a T_m_ of 55°C. This elevated T_m_ imparts several favorable characteristics on the liposome, including enhanced *in vivo* stability and reduced clearance rates compared to natural lipids^41^.

We synthesized liposomes via a modified form of our previously published SPIN method (**Figure 1a**)^42^, and concentrated using filter centrifugation, which we found was significantly superior to ultracentrifugation (**Figure 1b and 1c**). Liposomes display high stability when incubated at the T_m_ in up to 10% DMSO (as a vehicle of lipophilic molecules). We found that above this temperature, and when incubated with isopropyl or ethyl alcohol, liposomes rapidly fell apart resulting in the precipitation of lipids. However, previous reports by Balley et. al. have demonstrated that DSPC liposomes were stable for up to one hour at 70°C in up to 30% ethanol^43^. This discrepancy may be due to the increased levels of cholesterol (45%) in their liposomes as compared to 33% cholesterol in our experiments, which is reported to be the most stable^41,44,45^. Cholesterol has a stabilizing effect, creating order in the hydrophobic region at increased temperatures that could stabilize liposomes at these increased temperatures^46–47^. In our simulations, we found no interaction between the DXR and cholesterol (**Supplementary Figure 5**), but simulations to determine the effect of increased cholesterol on conformational energy may explain this discrepancy. In our simulated leaflet containing 33% cholesterol, we found little difference in the interactions between DXR and DSPC once the temperature was >T_m_. Electrostatic, VdW, and hydrogen bonding interactions (**Figures 4a and 4b**) were similar at both 55°C and 65°C. However, there was a larger than expected increase in conformational energy when comparing 55°C and 65°C membrane simulations. This increase in conformational energy also affects other membrane dynamics which can lead to loss of stability, such as the solvent accessible surface area (**Supplementary Figure 6**).

We showed equilibration as a robust approach to create potent targeting liposomes. The liposomes quickly equilibrated within an hour (**Figure 3b**) and can provide a targeted therapeutic effect (**Figure 3e**). Interestingly, we found that even at lower temperatures (i.e. <T_m_), DXR associates with liposomes. However, an increase in encapsulation was observed at higher temperatures, which is likely a direct result of the increased permeability of the membrane. The addition of cholesterol limits the phase transition of liposome membrane and maintains a liquid ordered phase^48^. Using NAMD to simulate the membrane dynamics and interatomic interactions, we investigated the association between liposomes and DXR at various temperatures. We found that the slight positive charge found on the amine group of DXR was highly attracted to the negative phosphate groups on lipids. This interaction elegantly explains the proficiency of active methods, wherein ion gradients (typically made from high intramolecular levels of sulphate and phosphate) attract slightly positive charged compounds such as DXR. The phosphate-NH_2_ attraction occurs through both long range (VdW) and intermediate range (electrostatic) interactions and allows for adsorption of DXR to the membrane. The adsorbed molecules are retained through all wash cycles but are quickly released once introduced to physiological conditions, as evidenced by the initial burst seen in **Figure 3d**. The concentration of DXR associated with liposomes decreases by 20% within the first two hours, but only 5% was released over the following 46 hours. This opens the exciting possibility of a tri-modal release, wherein liposomes can be engineered to contain DXR adsorbed to the liposome membrane (through low temperature equilibration) as well as encapsulated within the core (through passive encapsulation) and membrane (through high temperature equilibration).

In addition to drug compartmentalization, thermal equilibration also conveniently allows for the co-loading of compounds. Liposomes are promising avenues of increasing therapeutic index through combinatorial simultaneous delivery^49,50^. Recently, researchers have investigated the use of liposomes co-loaded with doxorubicin and paclitaxel, molecules that differ greatly in polarity^51^. Combined with our liposome synthesis and concentration strategy, we demonstrate here that the thermal equilibration approach effectively sequesters both hydrophilic and lipophilic compounds within targeting liposomes (**Figure 5b**) without requiring specialized steps. Furthermore, we can encapsulate both types of molecules at the same time, generating co-loaded targeting liposomes in a more efficient manner than passive encapsulation. While the goal of this work was to demonstrate that antibody coupling is possible during equilibration, future work characterizing the antibody insertion and efficiency (including maleimide hydrolysis) warrants further investigation. This will be highly variable depending on the intended application, and thus is not within the scope of the current work. Exploration on the compatibility of liposomes equilibrated in solvents other than DMSO, such as surfactants and emulsifiers, will greatly advance the use of equilibration in generating highly potent, co-loaded liposomes.

## Conclusions

In this work, we describe a novel strategy for rapidly generating potent targeting liposomes using the thermal equilibration approach. We characterize the stability and encapsulation potential of liposomes both experimentally and through molecular dynamic simulations. We then demonstrate the therapeutic capability by introducing targeted LDXR to cancer cells *in vitro* and evaluating cell viability. Finally, we demonstrate the universal nature of this approach by co-loading both hydrophilic and lipophilic molecules. Future work will consist of drug specific optimizations using microscopy and further cellular uptake assays to establish potent delivery systems. We believe that thermal equilibration offers unparalleled advantages compared to conventional encapsulation techniques with a variety of potential applications in the therapeutic and theranostic landscape.

## Author Information

### Author Contributions

NA and SR conceived the technology and designed the research plan. SR and CL synthesized liposomes and performed characterization experiments and data analysis. SS performed membrane simulations. SR wrote the initial manuscript draft. All authors discussed the results and edited the manuscript. All authors assume responsibility for their respective contributions and the overall work presented in the manuscript.

## Acknowledgments

The authors are grateful to Dr. Rohan Fernandes at The George Washington University for allowing the use of their analytical equipment and other resources for this project.

