## Supplementary Information for "Versatile Encapsulation and Synthesis of Potent Therapeutic Liposomes by Thermal Equilibration"

**Simulation System**

Simulations were performed on a lipid membrane system consisting of DSPC and cholesterol molecules in a 2:1 molar ratio respectively. Since all-atom MD simulations are computationally demanding, a section of the liposomal membrane was used to represent the thermodynamic changes observed at a local point in a liposome that allows the passive diffusion of small molecules like drugs. Charmm GUI[1,2] *Membrane Builder* was used to build the membrane system. The lipid bilayer was built in a rectangular box containing a total of 256 lipid molecules (128 on each layer of the bilayer known as a leaflet) and 128 cholesterol molecules (64 on each leaflet). Additionally, 30 water molecules per lipid molecule were present to ensure hydration of the lipid head. The effect of different temperatures on different lipid molecules (like DLPC, DMPC, and DPPC) has been previously studied by Zhuang *et. al*[3], using a bilayer of 72 lipid molecules (36 on each leaflet). The number of lipid molecules was increased by ~3.5 times for our simulations to build a bigger model representing the liposome. Furthermore, a bilayer system containing 256 molecules of DSPC has been used by Magarkar *et al*[4] who studied a molecular dynamics model of the liposome surface in the bloodstream. The $L_x$ and $L_y$ components of the bilayer are 10.5 nm each with an $L_z$ component of 4.25 nm. The total number of molecules in the system is given in Table 1.

**Supplementary Table 1**: Molecular Dynamic Simulation system information

| DSPC | Cholesterol | Water | Simulation temperatures (C) | Simulation time (ns) |
|---|---|---|---|---|
| 256 | 128 | 5534 | 37, 55, 65 | 100 |

**Molecular Dynamics simulation setup:**

The simulation used a standard TIP3 water model[5,6] and CHARMM36[7] lipid parameter set combined with the CHARMM parameters for the general force field of small molecules (CGenff)[8]. A neutralizing salt (NaCl) concentration of 0.05 mol/L was added to the system[9]. Periodic boundary conditions were applied in all three dimensions. Van der Waals interactions were switched off between 10 and 12 Å by a force based switching function[10]. In each simulation, Particle Mesh Edwald (PME)[11] was used for electrostatic interactions. NAMD[12,13] (Nanoscale Molecular Dynamics) was used to thermally equilibrate the bilayer system for 4 ns. RATTLE algorithm[14] was used to constrain all bond lengths involving hydrogen atoms.

All thermally equilibrated systems were used for simulation runs at different temperatures (Table 1). The simulation time step was 1fs and the data was collected every 20 ps. A simulation for each temperature was run for 100 ns. NPT ensemble was used for simulations with the pressure at 1 bar maintained by Nosè-Hoover Langevin[15] piston method. A constant temperature in each simulation was maintained by Langevin thermostat. The trajectories were visualized to create snapshots of the bilayer using Visual Molecular Dynamics (VMD).

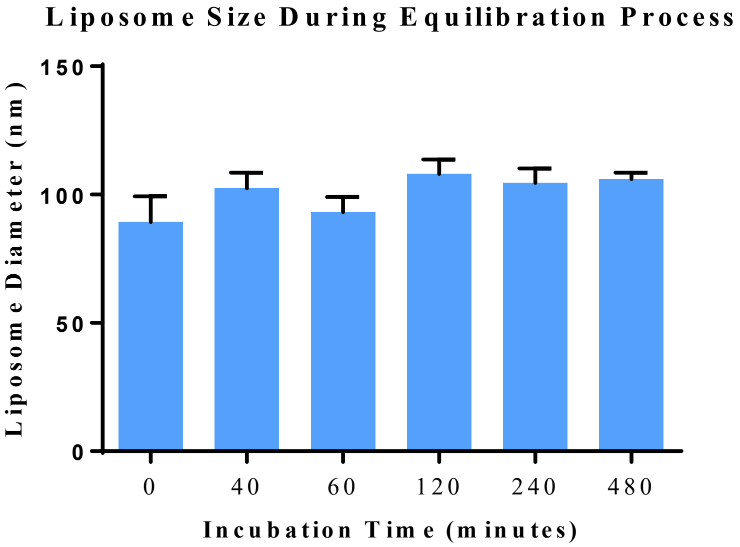


**Supplementary Figure 1**: Diameter of Liposomal DXR following $E_T$. The diameter of liposomes following increasing equilibration durations at 55 $^{O}$C was analyzed via DLS. Similar to Figure 2, the distribution of liposome diameter shows no significant changes over 480 minutes. This demonstrates the stability of the liposomes and is also used to evaluate the maximum equilibration concentration.

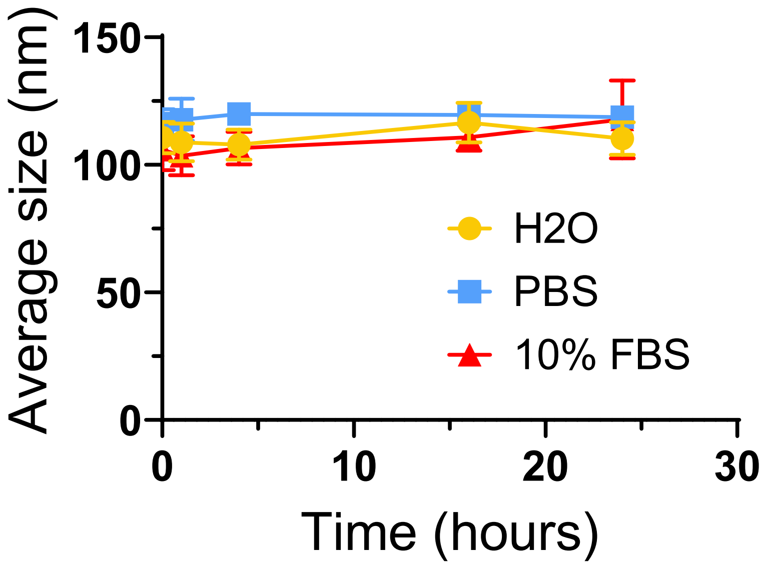


**Supplementary Figure 2**: The stability of liposomes in serum was tested. To comparison, the stability in $H_2O$ and PBS was also tested. The particles were dispersed in either $H_2O$, PBS, or 10% FBS in PBS and incubated at 37 ºC, and the particle size was recorded at each time point (0, 1, 4, 16, and 24 hours). The particle size change was negligible when the liposomes were incubated in $H_2O$ or PBS up to 24 hours 0.003 % and 2.11% respectably. When the particles were mixed with 10% FBS, the particle size was increased from 105.9 ± 10.2 nm to 117.8 ± 15.3 nm due to the protein absorption on the particle surface.

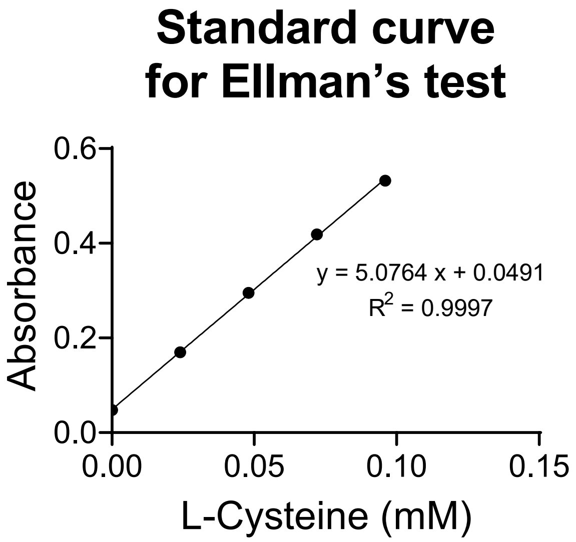


| | Average | std |
|---|---|---|
| Absorbance | 0.092 | 0.011 |
| DSPE-PEG-MAL (mM) | 0.059 | 0.003 |
| DSPE-PEG-MAL/DSPC ratio | 0.009 | 0.001 |

**Supplementary Figure 3**: Quantification of Maleimide:Lipid ratio was evaluated by Ellman's test. The result showed that the maleimide to DSPC lipid ratio is 0.009 which is less but close to its initial value of 0.01. Since the simulation showed one liposome consists of 256 lipids and 128 cholesterol, about 2-3 DSPE-PEG-MAL is inserted into each liposome.

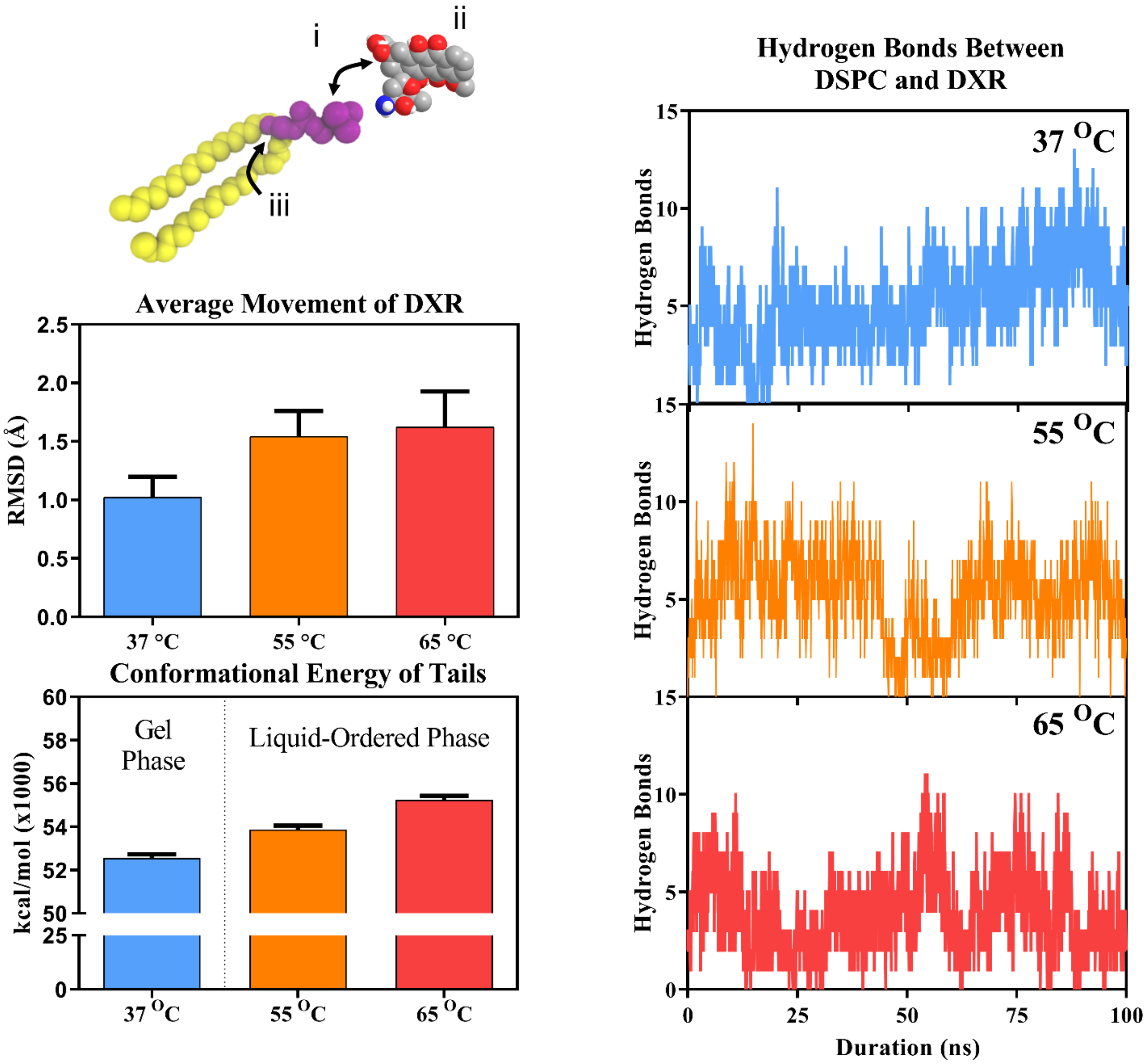


**Supplementary Figure 4**: Effects of increased temperature on DXR equilibration. (A) The stability of the DXR-lipid interactions is dictated by several factors including (i) hydrogen bonds, (ii) Brownian motion and diffusion of the DXR, and (iii) conformational energy of the tails which allows for diffusion through the membrane. (B) The DXR can be stabilized both inside and outside the liposome through hydrogen bonding between the oxygen and nitrogen atoms on the DXR and phosphates on the DSPC. (C) This stabilization, along with VdW and electrostatic interactions can have a profound effect on the net movement of the DXR through the liposome. (C) While temperature clearly has an effect on the fluidity of the membrane, it also plays a role in decreasing the liposome stability. Both the transition from 37 °C to 55 °C and from 55 °C to 65 °C result in an increase in conformational energy of 2.5%.

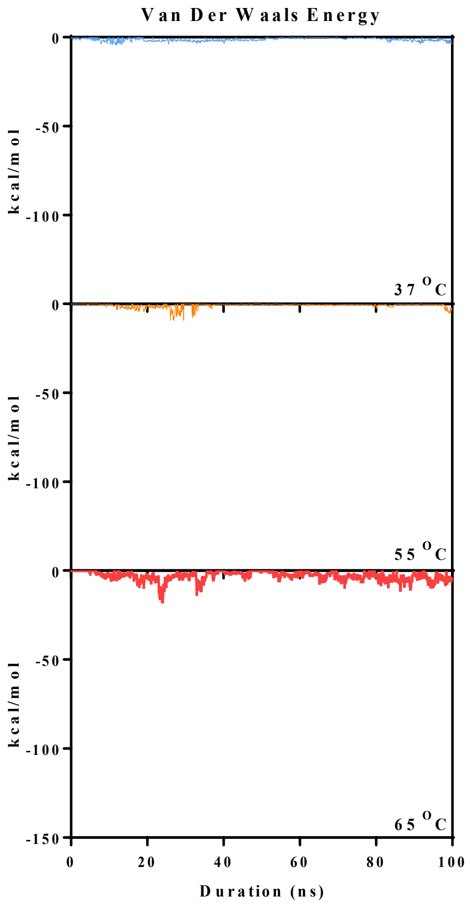


**Supplementary Figure 5**: Long-range VdW interactions between DXR and Cholesterol. VdW forces between cholesterol and DXR are minimal. At 65 $^{O}$C, lipid tails are characterized by higher motion exposing cholesterol molecules to DXR accounting for slight interactions between DXR and cholesterol as seen in the figure above. Below $T_m$, thermal movement of tails is limited and DXR is bound to lipid heads with no interactions with cholesterol. At $T_{m,}$ motions of tails are fluid but ordered which does not favor interactions between DXR and cholesterol. Cholesterol provides structural rigidity to liposomes. A higher interaction between DXR and cholesterol may affect the structural function of cholesterol and aid in the rapid breakdown of liposomes.

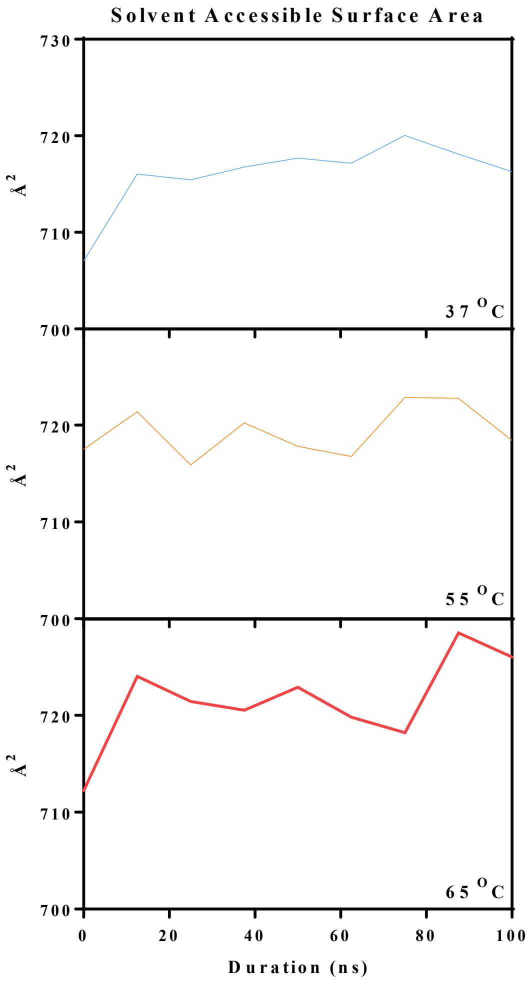


**Supplementary Figure 6**: Solvent Accessible Surface Area (SASA) of lipids at increasing temperatures analyzed at intervals of 10 ns. With increasing temperatures, SASA values showed a steady rise. A higher area of lipids is exposed to water due to fluctuations of the tails. The increased water exposed area with higher temperatures is another factor that contributes to the destabilization of the membrane. As the temperatures increased, SASA increased and could be a reason for the increased DXR movement into the lipid membrane during the simulation as well as destabilization of the membrane at higher temperatures.
